# Dissociable frequency regimes in human temporal cortex integrate facial and acoustic cues during natural speech

**DOI:** 10.64898/2026.03.02.709171

**Authors:** Jiawei Li, Kaixuan Bian, Xiaotao Hao, Youkun Qian, Jinsong Wu, Junfeng Lu, Yuanning Li

## Abstract

Face-to-face communication relies on the seamless integration of visual and acoustic cues, yet the spatiotemporal principles governing how the human brain dynamically represents and combines these multisensory streams remain largely unresolved. To address this, we recorded high-density electrocorticography (ECoG) from eight participants perceiving matched audiovisual, audio-only, and video-only continuous natural Mandarin speech. Using time–frequency–resolved encoding models, we reveal complementary, frequency-dependent integration regimes across the temporal lobe. We show that the superior temporal gyrus (STG) implements a feature-selective, auditory-dominant strategy, utilizing visual input to selectively strengthen low-frequency representations of lip-reading kinematics. Conversely, the middle temporal gyrus (MTG) acts as a higher-order multisensory hub, employing a frequency-selective strategy to broadly integrate diverse facial and articulatory features. Crucially, we demonstrate that access to visual information during perception significantly improves the acoustic and lexical accuracy of neural speech decoding and re-synthesis, with the MTG driving the largest gains in linguistic intelligibility. These findings uncover the dissociable neural architectures supporting robust multisensory perception, providing critical mechanistic insights for the development of next-generation, multimodal brain-computer interfaces.

## Introduction

Speech perception in real-world settings is fundamentally multisensory computation [1–3]. In face-to-face communication, the brain must dynamically synthesize not only acoustic cues (spectrotemporal structure, phonetic content, prosody) but also visual signals that are tightly coupled to articulation and social intent, including lip kinematics, jaw dynamics, and broader facial actions (e.g., eyebrow movements, blinks, affective expressions[4–6]). These visual cues improve intelligibility, especially under acoustic ambiguity or noise, and they can shape the perceived identity, timing, and meaning of speech. Therefore, understanding how the brain integrates these distinct streams is central to a complete model of speech perception. Importantly, recent progress has begun to localize critical circuitry for audiovisual integration to posterior superior temporal cortex and adjacent temporal-lobe regions[7–11], establishing plausible anatomical substrates for cross-modal interactions during speech. Yet, the precise spatiotemporal coding principles governing how the human brain represents and integrates auditory and facial cues across cortical areas and frequency bands during continuous, natural speech remain unclear.

Over the past decade, substantial progress has localized critical circuitry for speech and face processing within the temporal lobe [12, 13]. A major emphasis has been on high-frequency activity, particularly high gamma band (70-150 Hz), in superior temporal gyrus and sulcus, which reliably tracks natural speech and reveals rich population-level spatiotemporal structure encoding spectrotemporal acoustics, phonetic features, and higher-order linguistic variables[14–18]. In parallel, a complementary line of work has shown that lower-frequency oscillations in the superior temporal gyrus (STG), especially in the theta range, align with temporal and linguistic structure in continuous speech, consistent with roles in segmentation and multi-timescale tracking of speech dynamics[19]. Encoding-model approaches, including temporal receptive field frameworks[20], have provided quantitative descriptions of these mappings and clarified how speech representations unfold over time within STG populations.

In parallel, studies of face perception and facial motion have implicated temporal-lobe regions often associated with the visual and face-processing streams. Beyond relatively static face identity signals in the ventral pathway, converging evidence suggests that a more dorsal temporal route, including regions such as the middle temporal gyrus (MTG) and the superior temporal sulcus (STS), is particularly engaged by dynamic facial cues that unfold over time, such as mouth movements and broader facial actions [21–24]. Importantly, these facial-motion representations can be expressed across frequencies, with several reports highlighting reliable structure in lower-frequency activity[24, 25] (below 25 Hz) alongside high-frequency responses (high-gamma range)[26]. Together, these observations raise the possibility that audiovisual speech engages partly distinct frequency regimes across temporal cortex, with high-gamma responses strongly characterizing auditory speech coding in STG, and lower-frequency dynamics contributing substantially to the representation of dynamic facial cues in temporal regions associated with face processing.

However, these two streams of research have largely evolved in isolation. Most audiovisual speech studies have focused on how visual cues influence neural dynamics within the core auditory speech network, particularly STG and STS[27, 28], often through the lens of high-gamma activity or condition-contrast responses[29–31]. By comparison, it remains less clear how temporal regions traditionally studied for visual facial processing incorporate auditory speech information, and whether auditory input modulates their facial representations, including those expressed in lower-frequency bands. More broadly, the field still lacks a feature-resolved, frequency-resolved account of audiovisual integration that specifies where integration occurs across temporal-cortical areas, and which frequency bands carry modality-specific versus multisensory representations during continuous, natural speech.

To address this, we recorded high-density electrocorticography (ECoG) from eight human participants during the perception of matched audiovisual, audio-only, and video-only continuous, natural speech. We hypothesized that audiovisual integration is not a monolithic process, but rather a regionally specialized and frequency-dependent computation. Specifically, we predicted a functional dissociation: the STG would preferentially integrate speech-critical visual cues (e.g., lip kinematics) via high-frequency, local feature encoding, whereas the MTG would serve as a broader multisensory hub, integrating diverse facial and auditory features through temporally structured, lower-frequency bands.

We tested this architecture using a comprehensive, computationally rigorous framework. First, we constructed time–frequency–resolved encoding models using precise visual (Facial Action Units) and articulatory (Articulatory Kinematic Trajectories) features derived directly from natural speech stimuli. Second, we directly linked these representational principles to functional outcomes by utilizing the recorded neural activity to decode and re-synthesize naturalistic speech. By quantifying the regional and spectral gains afforded by multisensory input, this framework provides a definitive, frequency-resolved account of how the human temporal cortex bridges facial and acoustic dynamics to achieve robust speech perception.

## Results

### Experiment overview

This study investigates the dynamic neural encoding of audiovisual speech perception using naturalistic, ecologically valid stimuli. Participants (n = 8) were presented with matched audiovisual (AV), audio-only (A), and video-only (V) segments of continuous natural Mandarin speech drawn from professional news broadcasts, containing continuous, fluent natural speech and clear articulatory movements (**Fig. 1A**). ECoG recordings were obtained from these patients during the experiments. Electrodes were primarily placed over the STG and MTG. As shown in **Fig. 1B**, individual participants’ electrode coverage was widely distributed across these areas, with distinct spatial patterns reflecting inter-individual variability in surgical targeting. Collectively, the dataset included 1408 electrodes covering STG and MTG (**Fig. 1C**), allowing for high-resolution mapping of neural activity across spatial, temporal and frequency domains.

**Fig. 1.**
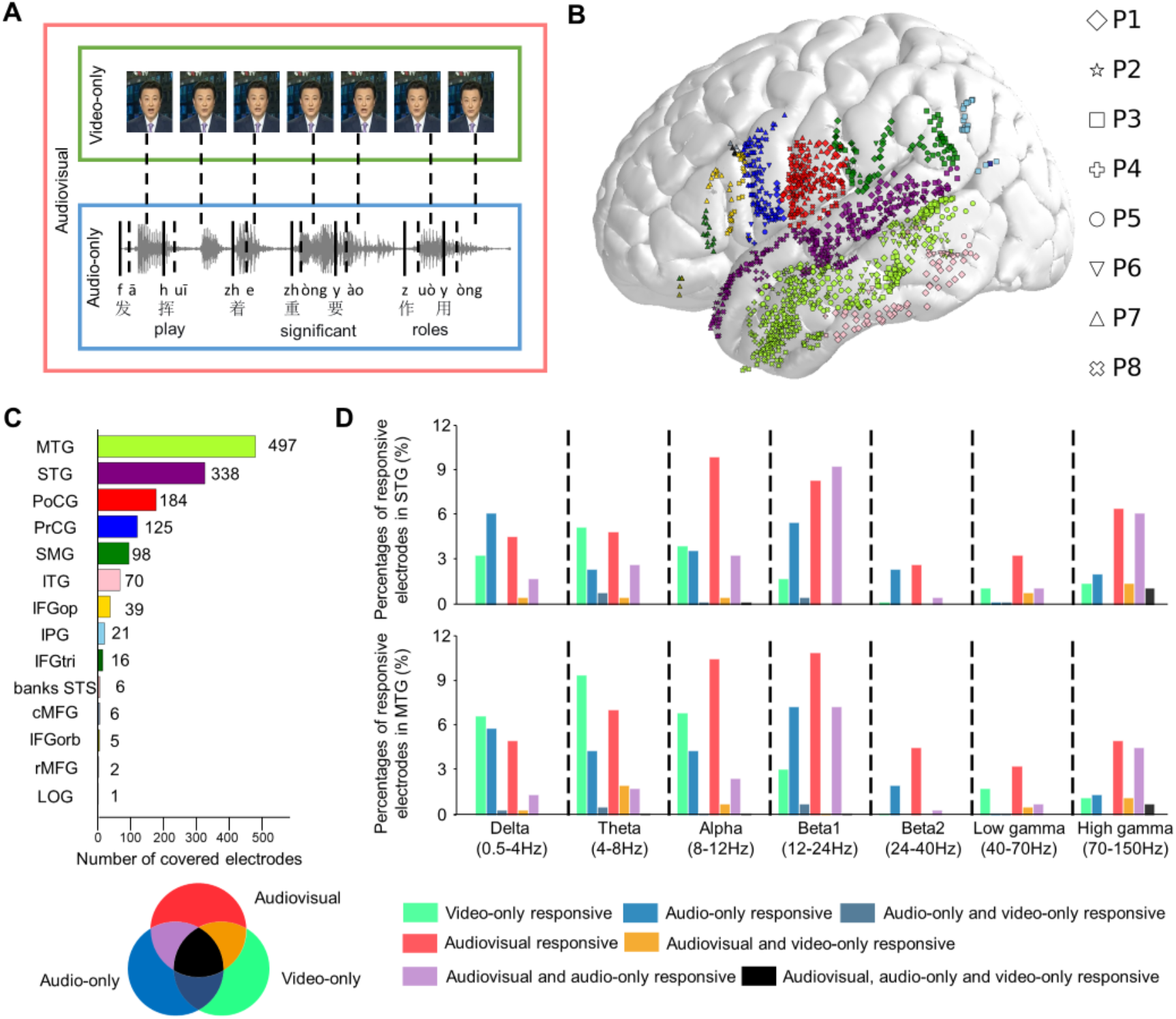
Experimental design and electrophysiological coverage. **A**. Overview of stimuli presented to participants under three modalities: audiovisual (AV), audio-only (A), and video-only (V) conditions. The audio component consisted of natural continuous Mandarin speech delivered by a professional broadcaster, illustrated here with a demonstrative waveform segment of the phrase “发挥着重要作⽤” (“play significant roles”), annotated with syllables and phonetic components (vowels and consonants). The video component included high-resolution clips showing clear articulatory movements during speech production. **B**. Electrode coverage across eight participants (P1–P8), mapped onto a standard cortical surface model. Different brain regions are color-coded, and individual participants are distinguished by unique marker shapes. **C**. Total number of electrodes covering each brain region across all participants. Colors correspond to those in panel **B**. **D**. Percentages of responsive electrodes in the superior temporal gyrus (STG) and middle temporal gyrus (MTG) across frequency bands (Delta: 0.5–4 Hz; Theta: 4–8 Hz; Alpha: 8–12 Hz; Beta1: 12– 24 Hz; Beta2: 24–40 Hz; Low Gamma: 40–70 Hz; High Gamma: 70–150 Hz). Response types are categorized as: video-only responsive (green), audio-only responsive (blue), audio-only and video-only responsive (blue-black), audiovisual responsive (red), audiovisual and video-only responsive (orange), audiovisual and audio-only responsive (purple), and responsive to all three modalities (black). Dashed lines separate distinct frequency bands.

We addressed three key questions: First, we examined how audiovisual integration distributed over different cortical areas and frequency bands. Second, we investigated how event-related potentials (ERPs) differ under AV, A, and V conditions, assessing the extent to which each modality contributes uniquely to the neural response. Third, we explore how specific visual features and articulatory-phonetic features, such as facial Action Units (AUs) and Articulatory Kinematic Trajectories (AKTs), are modulated by stimuli modality, and quantify their predictive power on neural activity. Specifically, we analyze whether the introduction of audio or visual information in AV conditions enhances the encoding of auditory or visual features compared to unimodal conditions. To answer these questions, we examined the distribution of responsive electrodes across the full spectrum (0.5– 150 Hz) by comparing event-related potentials to baseline resting-state potentials. Our analysis reveals widespread responsiveness in both STG and MTG to speech stimuli across multiple frequency bands (**Fig. 1D**). In STG, the proportion of electrodes showing significant responses varied by band: delta (16.0%), theta (16.3%), alpha (21.3%), beta1 (24.6%), beta2 (5.9%), low gamma (7.1%), and high gamma (18.3%). In MTG, the corresponding proportions were delta (19.1%), theta (24.7%), alpha (24.5%), beta1 (28.8%), beta2 (7.0%), low gamma (6.8%), and high gamma (13.9%). Notably, a substantial subset of electrodes exhibited multimodal sensitivity, responding to both auditory and visual cues (i.e., to at least two of the three stimulus types: A-only, V-only, or AV). Across frequency bands, the average proportion of multimodal electrodes was 4.7% in STG and 3.9% in MTG, with the highest proportions observed in the beta1 (STG: 9.5%; MTG: 8.2%) and high gamma bands (STG: 8.6%; MTG: 6.4%). These findings suggest functional convergence in these temporal regions during speech perception.

### Distinct modality sensitivity across temporal cortex and frequency bands

To characterize the neural substrates of audiovisual speech processing, we first mapped the spatial distribution of electrodes showing significant modulation by audiovisual (AV), audio-only (A), or video-only (V) stimuli across frequency bands. Responsive electrodes were defined as those showing task-related ERP amplitudes significantly different from rest-state baselines (p < 0.05, two-sided, Bonferroni corrected). Density of responsive electrodes can be observed throughout the delta (0.5–4 Hz) to high gamma (70–150 Hz) range revealed by kernel density estimation (**Fig. 2A**). Many electrodes responded to multiple conditions, such as both A and V, or exclusively to AV, indicating heterogeneous modality sensitivity within the human brain, especially the temporal lobe (e.g. E1– E7; **Fig. 2B**). However, no systematic similarity between AV, A, and V conditions was observed at this stage.

**Fig. 2.**
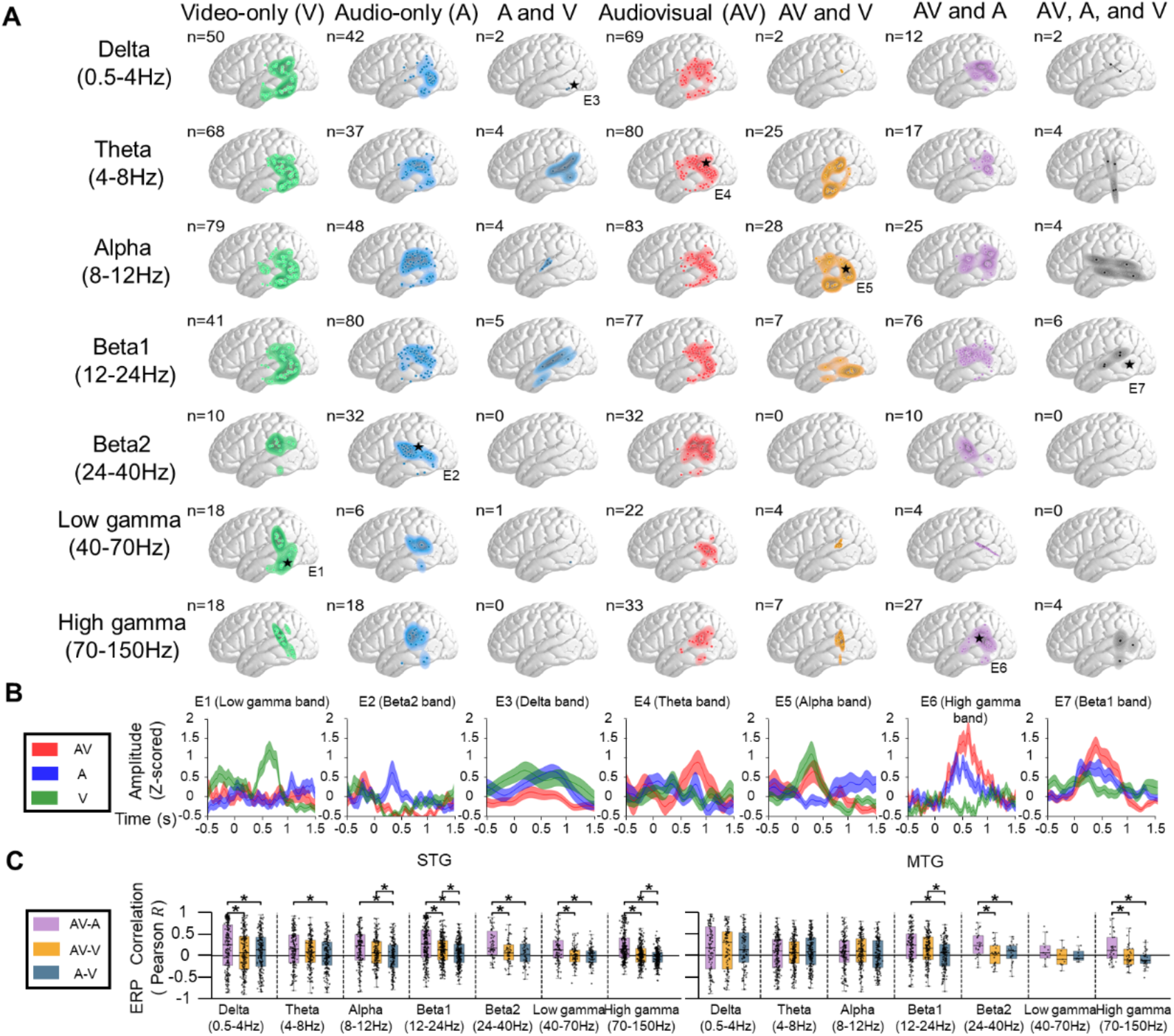
Distribution of responsive electrodes, representative event-related potentials (ERPs), and cross-modal ERP correlations. **A**. Spatial distribution of electrodes responsive to different stimulus conditions (audiovisual [AV], audio-only [A], video-only [V]) across frequency bands (Delta: 0.5–4 Hz; Theta: 4–8 Hz; Alpha: 8– 12 Hz; Beta1: 12–24 Hz; Beta2: 24–40 Hz; Low gamma: 40–70 Hz; High gamma: 70–150 Hz), visualized on cortical surface models. Each column represents a unique response pattern (e.g., AV-only, A and V, etc.), with the total number of responsive electrodes indicated (n). Kernel density estimation (KDE) maps show the spatial concentration of responses. Demonstrative electrodes (E1– E7) are marked in each panel for subsequent analysis. **B**. Average event-related potentials (ERPs) from the seven demonstrative electrodes (E1–E7), time-locked to sentence onset (t = 0 s). ERPs are z-scored and plotted separately for AV (red), A (blue), and V (green) conditions. The x-axis indicates time relative to stimulus onset, and the y-axis shows z-scored amplitude. **C**. Pearson correlation coefficients between pairwise ERP patterns (AV vs. A, AV vs. V, A vs. V) in the superior temporal gyrus (STG) and middle temporal gyrus (MTG), computed across all electrodes responsive to at least one condition in each frequency band. Correlations are shown as boxplots with individual data points. Asterisks (*) denote statistically significant correlations (p < 0.05), indicating that the ERP patterns for two stimulus types are more strongly correlated than the other two comparisons within a given band and region.

### Multisensory neural responses reveal auditory dominance in STG and frequency-tuned integration in MTG

We next asked whether the temporal response structure in AV resembled A or V more strongly, and whether this depended on region and frequency. To quantify similarity at scale, we computed pairwise R^2^ between ERP waveforms across electrodes for each band (Fig. 2C). In STG, AV responses were consistently more similar to A than to V across most frequency bands, indicating an auditory-dominant response regime under multisensory input (Fig. 2C: AV-A correlations vs. AV-V correlations: t(200) = 2.61, p = 9.62 × 10^−3^ in delta band, t(260) = 4.56, p = 7.95 × 10^−6^ in beta1 band, t(61) = 3.51, p = 8.50 × 10^−4^ in beta2 band, t(94) = 3.82, p = 2.43 × 10^−4^ in low gamma band, t(274) = 7.39, p = 1.75 × 10^−12^ in high gamma band, paired two-sided t-test; AV-A correlations vs. A-V correlations: t(200) = 2.47, p = 1.44 × 10^−2^ in delta band, t(183) = 3.51, p = 5.62 × 10^−4^ in theta band, t(175) = 5.74, p = 4.13 × 10^−8^ in alpha band, t(260) = 8.46, p = 1.95 × 10^−15^ in beta1 band, t(61) = 4.44, p = 3.81 × 10^−5^ in beta2 band, t(94) = 3.87, p = 1.98 × 10^−4^ in low gamma band, t(274) = 10.52, p = 5.70 × 10^−22^ in high gamma band, paired two-sided t-test). In contrast, MTG showed frequency-dependent AV similarity patterns, with AV–A advantages emerging in more restricted bands (notably beta2 and gamma/high-gamma comparisons), consistent with a spectrally selective integration profile rather than a uniformly auditory-driven regime (Fig. 2C: AV-A correlations vs. AV-V correlations: t(27) = 2.88, p = 7.61 × 10^−3^ in beta2 band, t(34) = 5.01, p = 1.14 × 10^−2^ in high gamma band, paired two-sided t-test), while in beta (12-40 Hz) and high gamma (70-150Hz) bands compared with A-V correlations (AV-A correlations vs. A-V correlations: t(161) = 6.55, p = 7.28 × 10^−10^ in beta1 band, t(27) = 3.09, p = 4.57 × 10^−3^ in beta2 band, t(34) = 3.53, p = 1.20 × 10^−3^ in high gamma band, paired two-sided t-test).

These findings suggest a functional dissociation between STG and MTG: while STG maintains a stable, auditory-centric code for speech across sensory contexts, whereas MTG supports frequency-tuned integration of visual articulatory information, potentially enabling context-sensitive perceptual synthesis during natural communication.

### Temporal receptive field models reveal cross-band encoding of facial and articulatory dynamics

To move beyond condition contrasts and identify which features drive neural responses, we further investigated the extent to which specific features of facial and articulatory motion could predict cortical activity in the superior and middle temporal gyrus (STG and MTG). To this end, we introduced two key descriptors: the Facial Action Units (AUs, **Fig. 3A and 3B**) and the Articulatory Kinematic Trajectories (AKTs, **Fig. 3A and 3C**). AUs provide a standardized, objective description of facial dynamics by quantifying discrete facial muscle activations (see Methods). AKTs instead capture continuous, time-varying articulator motions during speech, reflecting coordinated, context-dependent kinematics underlying fluent production.

**Fig. 3.**
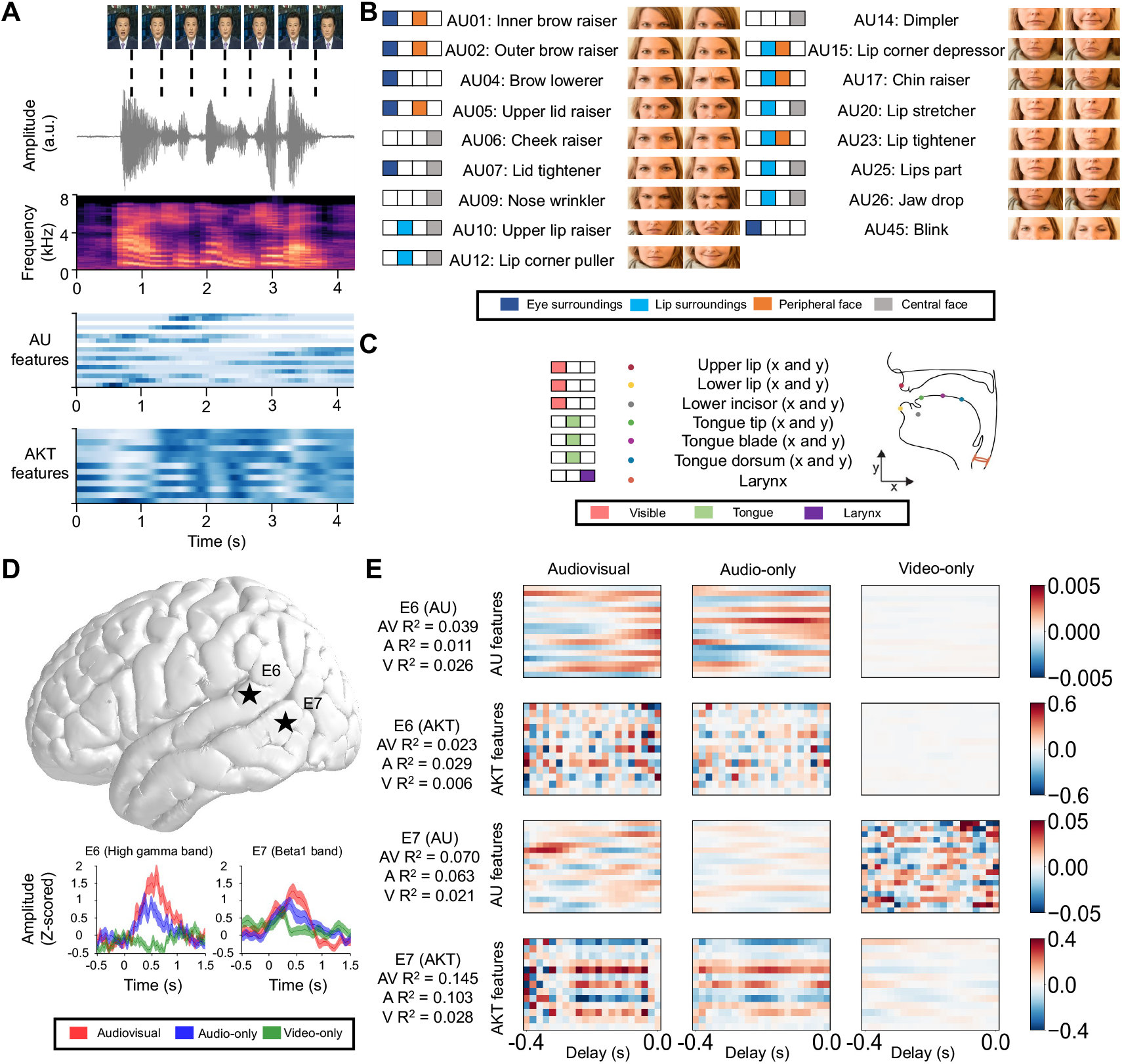
Facial Action Units (AUs), Articulatory Kinematic Trajectories (AKTs) and their neural correlates. **A**. Example of multimodal speech representation, showing a sequence from video clips of facial expressions, the corresponding audio waveform, mel-spectrogram, features of facial action units (from AU01, inner brow raiser, to AU45, lip corner puller), and Articulatory Kinetic Trajectories (AKT) features. **B**. Schematic representation of facial Action Units (AUs) and their spatial distribution on the face, with visual illustrations showing muscle activation patterns for each AU (e.g., AU01: inner brow raiser; AU45: blink). Each AU is represented by two images depicting its activation in neutral and activated states. AUs are grouped into four anatomical categories based on facial location: Eye surroundings (blue), Lip surroundings (cyan), Peripheral face (orange), and Central face (gray), as indicated in the legend. **C**. Schematic representation of the Articulatory Kinematic Trajectories (AKT) features and their anatomical locations within the human oral cavity. Each AKT feature (except larynx) is two-dimensional, capturing motion along the x- and y-axes. Features are grouped into three categories based on visibility and anatomical function: Visible (upper lip, lower lip, lower incisor), Tongue (tongue tip, tongue blade, tongue dorsum), and Larynx. The legend indicates the color coding for each group. **D**. Electrode locations and the average event-related potentials (ERPs) from the two demonstrative electrodes (E6 and E7), time-locked to sentence onset (t = 0 s). ERPs are z-scored and plotted separately for AV (red), A (blue), and V (green) conditions. The x-axis indicates time relative to stimulus onset, and the y-axis shows z-scored amplitude. **E**. Temporal receptive fields (regression weights) for electrodes 6 and 7, mapping AU and AKT features to neural responses under audiovisual, audio-only, and video-only stimulation conditions. For electrode 6, regression weights for both AU and AKT features are shown; for electrode 7, only AU feature weights are displayed.

To evaluate the predictive power of these features, we fit temporal receptive field (TRF) models under audiovisual (AV), audio-only (A), and video-only (V) conditions and quantified predictive performance (full-model R^2^) across frequency bands for STG and MTG.

For AUs, which primarily reflect visual facial muscle movements, we compared AV and A conditions to evaluate the contribution of visual input. In the STG, AU-related encoding showed comparable predictive power across modalities except alpha (8–12 Hz) band (**Fig. 4A**, t(116) = 3.48, p = 7.16 × 10^−4^ in alpha band, two-sided t-test). In contrast, the MTG exhibited significantly higher AU-based R^2^ values in AV than in A or V, particularly in beta (12–40 Hz) bands (**Fig. 4B**, AV > A: t(182) = 3.80, p = 1.95 × 10^−4^ in beta1 band, two-sided t-test; AV > V: t(182) = 4.13, p = 5.91 × 10^−5^ in beta1 band, t(34) = 2.84, p = 2.20 × 10^−3^ in beta2 band, two-sided t-test). This suggests that MTG integrates visual facial features more effectively when combined with auditory input, supporting its role in holistic face perception and multimodal social signal processing.

**Fig. 4.**
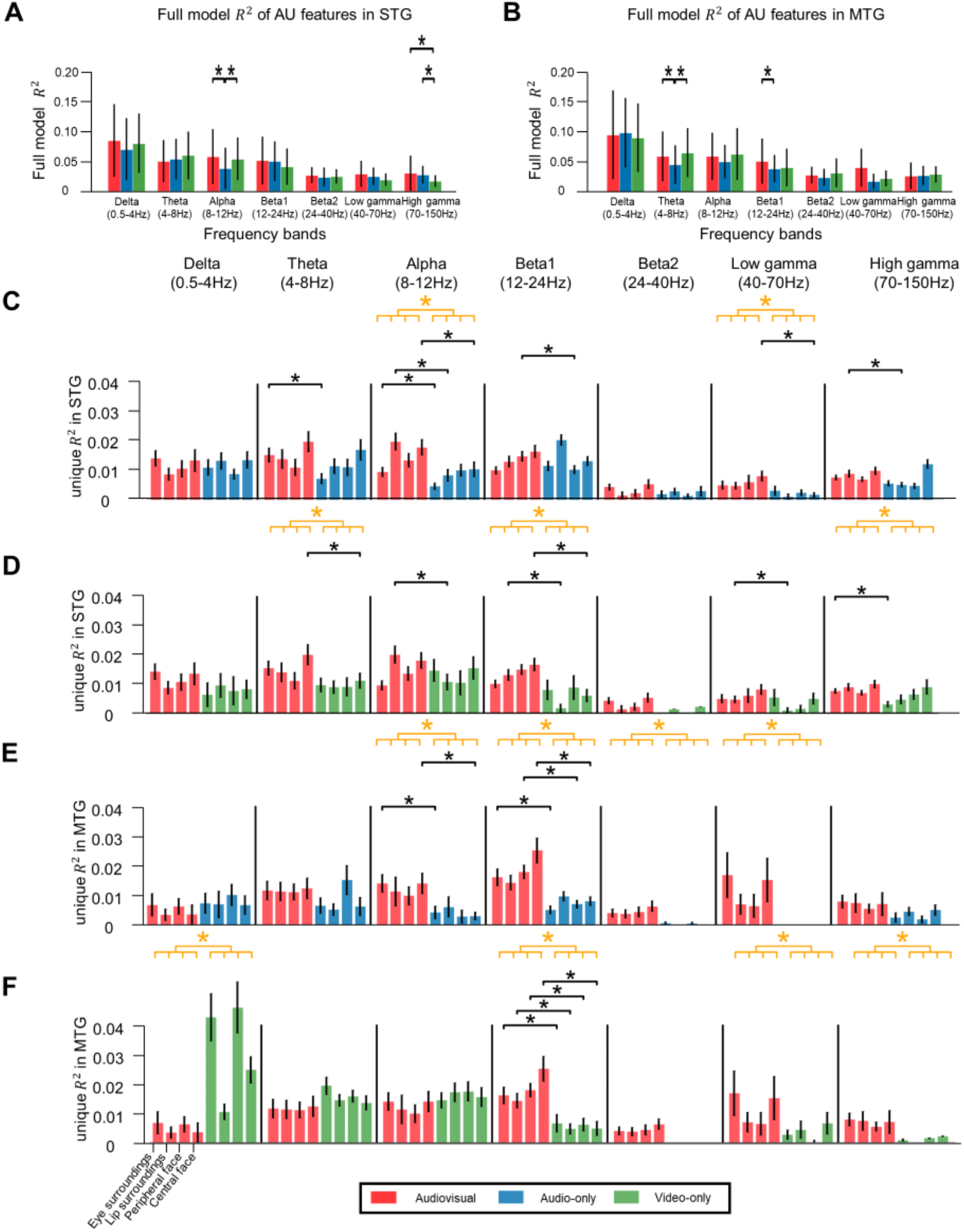
Unique neural predictions of Facial Action Units (AUs) across anatomical facial regions. **A**-**B**. Predictive power (measured by full model R^2^) of and Facial Action Units (AUs) in explaining brain activity in the superior temporal gyrus (STG; panels **A**) and middle temporal gyrus (MTG; panels **B**), across frequency bands (Delta: 0.5–4 Hz; Theta: 4–8 Hz; Alpha: 8–12 Hz; Beta1: 12–24 Hz; Beta2: 24–40 Hz; Low gamma: 40–70 Hz; High gamma: 70–150 Hz). Results are shown for three stimulus conditions: audiovisual (AV, red), audio-only (A, blue), and video-only (V, green). Data were modeled using Temporal Response Functions (TRF), with R^2^ indicating the proportion of variance in neural activity explained by each feature. Asterisks (*) denote statistically significant effects (p < 0.05), comparing the predictive performance of AKT or AUs across conditions and frequency bands. **C–F**. Unique R^2^ values (i.e., variance in neural activity uniquely explained by each AU group, after accounting for other predictors) in the superior temporal gyrus (STG; **B, C**) and middle temporal gyrus (MTG; **D**,**E**), across frequency bands (Delta: 0.5–4 Hz; Theta: 4–8 Hz; Alpha: 8–12 Hz; Beta1: 12–24 Hz; Beta2: 24–40 Hz; Low gamma: 40–70 Hz; High gamma: 70–150 Hz). Results are shown for audiovisual (AV, red), audio-only (A, blue), and video-only (V, green) conditions. Black bars indicate that the AV condition yields higher unique R^2^ than either unimodal condition within a given feature group and frequency band. Orange asterisks (*) denote significant differences (p < 0.05) from ANOVA tests, indicating that the overall unique R^2^ of the AV feature group exceeds that of individual modalities.

For AKTs, which primarily encode auditory-driven articulatory kinematics, we focused on comparing AV and V conditions to assess the contribution of auditory input. In the STG, significant predictive power was observed in the beta (12–40 Hz) and high gamma (70-150 Hz) bands, indicating enhanced auditory integration of articulatory motion (**Fig. 5A**, t(164) = 6.58, p = 6.04 × 10^−10^ in beta1 band, t(38) = 2.25, p = 0.30 × 10^−2^ in beta2 band, t(112) = 3.16, p = 2.01 × 10^−3^ in high gamma band, two-sided t-test). In MTG, AKT encoding was widespread from delta to beta1 bands (0.5-24 Hz) under AV conditions (**Fig. 5B**, t(80) = 3.38, p = 1.09 × 10^−3^ in delta band, t(118) = 2.36, p = 2.02 × 10^−2^ in theta band, t(94) = 3.36, p = 1.10 × 10^−3^ in alpha band, t(116) = 2.39, p = 1.82 × 10^−2^ in beta1 band, two-sided t-test). Notably, AV consistently outperformed V in these lower-to-mid frequency ranges, suggesting a broader role for MTG in integrating auditory and visual articulatory signals.

**Fig. 5.**
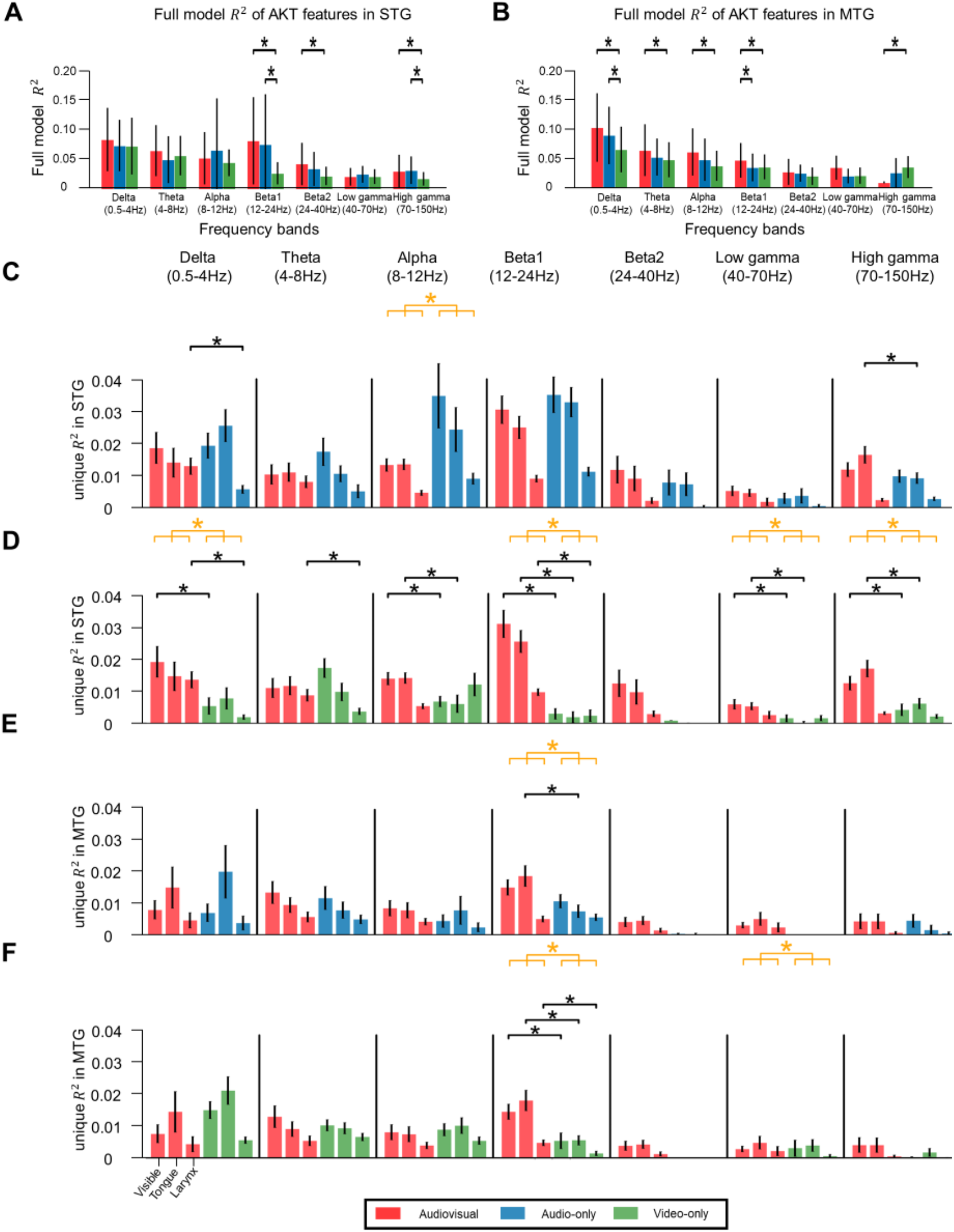
Unique neural predictions of Articulatory Kinematic Trajectories (AKT) across anatomical and functional groups. **A**-**B**. Predictive power (measured by full model R^2^) of and Articulatory Kinematic Trajectories (AKT) in explaining brain activity in the superior temporal gyrus (STG; panels **A**) and middle temporal gyrus (MTG; panels **B**), across frequency bands (Delta: 0.5–4 Hz; Theta: 4–8 Hz; Alpha: 8–12 Hz; Beta1: 12–24 Hz; Beta2: 24–40 Hz; Low gamma: 40–70 Hz; High gamma: 70–150 Hz). Results are shown for three stimulus conditions: audiovisual (AV, red), audio-only (A, blue), and video-only (V, green). Data were modeled using Temporal Response Functions (TRF), with R^2^ indicating the proportion of variance in neural activity explained by each feature. Asterisks (*) denote statistically significant effects (p < 0.05), comparing the predictive performance of AKT or AUs across conditions and frequency bands. **C–F**. Unique R^2^ values (i.e., variance in neural activity explained exclusively by each AKT feature group, after accounting for other predictors) in the superior temporal gyrus (STG; B) and middle temporal gyrus (MTG; C), across frequency bands (Delta: 0.5–4 Hz; Theta: 4–8 Hz; Alpha: 8–12 Hz; Beta1: 12–24 Hz; Beta2: 24–40 Hz; Low gamma: 40–70 Hz; High gamma: 70–150 Hz). Results are shown for audiovisual (AV, red), audio-only (A, blue), and video-only (V, green) conditions. Black bars indicate that the AV condition yields higher (p < 0.05) unique R^2^ than either unimodal condition within a given feature group and frequency band. Orange asterisks (*) denote significant differences (p < 0.05) from ANOVA tests, indicating that the overall unique R^2^ of the AV feature group exceeds that of individual modalities.

Together, these findings reveal a functional dissociation between STG and MTG: while STG prioritizes modality-specific integration of auditory-driven articulatory kinematics (via AKTs in beta1/beta2), MTG supports broader multimodal integration of both articulatory and facial expression features, particularly in mid-frequency bands.

### STG specifically selectively integrates visual lip-kinematics via frequency-wide encoding

In the STG, multisensory integration relative to unisensory conditions produces widespread enhancement across multiple frequency bands, but sharply constrained by feature type. When visual information is introduced (AV > A), only lip related facial action units show significant unique R^2^ gains, observed in the alpha band (t(46)=2.66, p=4.89×10^−3^) and high gamma band (t(50)=2.11, p=1.90×10^−2^), with additional modest effects in eye surroundings (alpha: t(46)=2.02, p=2.36×10^−2^; theta: t(27)=2.46, p=8.96×10^−3^) and central face features (alpha: t(46)=1.78, p=3.94×10^−2^; low gamma: t(18)=2.03, p=2.71×10^−2^; **Fig. 4C**). ANOVA confirmed significant main effects in alpha (p=3.62×10^−4^) and low gamma (p=1.62×10^−2^) bands. For articulatory kinematics, visual enhancement is minimal, with only laryngeal motion in delta (t(23)=2.57, p=6.77×10^−3^) and tongue in high gamma (t(50)=2.04, p=2.24×10^−2^) showing effects (**Fig. 5C**). By contrast, when auditory information is introduced (AV > V), STG exhibits robust, multi-band enhancement across nearly all articulatory kinematic features: visible, tongue, and larynx features gain significant unique R^2^ increases across delta, theta, alpha, beta1, low gamma, and high gamma bands (e.g., visible in delta: t(23)=1.99, p=2.72×10^−2^; visible in alpha: t(46)=2.11, p=1.96×10^−2^; tongue in beta1: t(57)=2.53, p=6.92×10^−3^; larynx in beta1: t(57)=2.53, p=6.99×10^−3^; tongue in low gamma: t(18)=2.94, p=3.60×10^−3^; **Fig. 5D**), with significant ANOVA effects in delta (p=1.92×10^−3^), beta1 (p=3.14×10^−4^), low gamma (p=5.13×10^−3^), and high gamma (p=4.74×10^−3^). Facial AU enhancement under auditory input remains focused on lip surroundings features (alpha: t(46)=1.78, p=4.03×10^−2^; beta1: t(57)=2.15, p=1.76×10^−2^) with additional effects in central face (theta: t(27)=1.89, p=3.26×10^−2^; beta1: t(57)=1.82, p=3.65×10^−2^; **Fig. 4D**), supported by significant ANOVA in theta (p=1.15×10^−2^), beta1 (p=3.37×10^−3^), and high gamma (p=3.39×10^−2^). Consistent with this auditory-driven broad enhancement, the example electrode E6 in STG (high gamma band) shows that R^2^ for articulatory kinematics under AV (0.023) is comparable to that under A (0.029) and both exceed V (0.006), with regression weights for AV closely matching those for A (**Fig. 3D–E**). This pattern reveals STG’s functional asymmetry: auditory input broadly optimizes articulatory representations across frequencies, while visual input selectively sharpens lip reading related features.

### Multisensory encoding in MTG exhibits feature-wide gains but concentrates in specific frequency bands

Unlike STG’s feature selectivity, MTG demonstrates frequency-specific integration that encompasses diverse feature categories. Under visual enhancement (AV > A), facial action units show widespread gains across multiple feature groups that are primarily concentrated in the beta1 band: eye surroundings (t(32)=3.06, p=1.70×10^−3^), peripheral face (t(32)=3.67, p=2.77×10^−4^), and central face (t(32)=3.32, p=7.92×10^−4^) all exhibit significant enhancement, with ANOVA confirming a highly robust beta1 effect (p=5.96×10^−8^; **Fig. 4E**). Modest additional contributions appear in the alpha band (eye surroundings: t(21)=2.04, p=2.54×10^−2^; central face: t(21)=2.04, p=2.55×10^−2^; ANOVA p=4.86×10^−3^) and low gamma band (ANOVA p=1.62×10^−2^). For articulatory kinematics, visual enhancement is highly circumscribed, with only tongue features showing gains in the beta1 band (t(32)=2.66, p=5.14×10^−3^; ANOVA p=1.24×10^−2^; **Fig. 5E**). Under auditory enhancement (AV > V), MTG’s integration becomes even more sharply focused on the beta1 band while expanding across features: all facial AU groups including eye surroundings (t(32)=1.86, p=3.49×10^−2^), lip surroundings (t(32)=2.26, p=1.45×10^−2^), peripheral face (t(32)=2.93, p=2.71×10^−3^), and central face (t(32)=2.85, p=3.32×10^−3^) show significant beta1 enhancement, with ANOVA yielding an extremely robust main effect (p=3.01×10^−6^; **Fig. 4F**). All articulatory kinematic features also gain beta1 enhancement (visible: t(32)=2.19, p=1.70×10^−2^; tongue: t(32)=2.37, p=1.11×10^−2^; larynx: t(32)=2.17, p=1.77×10^−2^; ANOVA p=8.79×10^−4^; **Fig. 5F**). Additional modest contributions appear in low gamma (ANOVA p=3.16×10^−2^) and high gamma (ANOVA p=2.18×10^−2^) bands for facial AUs. Supporting this bidirectional multisensory facilitation, the example electrode E7 in MTG (beta1 band) demonstrates that R^2^ for both facial AUs and articulatory kinematics under AV (AU: R^2^=0.070, AKT: R^2^=0.145) exceeds that under either A (AU: R^2^=0.063, AKT: R^2^=0.103) or V (AU: R^2^=0.021, AKT: R^2^=0.028) alone (**Fig. 3D–E**). Thus, MTG employs the beta1 band as its primary integration hub, supplemented by alpha and low gamma activity, to broadly encode both articulatory and expressive facial features.

### STG and MTG implement complementary encoding for multisensory integration

Together, these findings reveal specialized organizations in temporal cortex multisensory processing. STG adopts a frequency-wide but feature-focused coding strategy: auditory input drives comprehensive enhancement of articulatory kinematics across multiple frequency bands, while visual input selectively boosts lip reading related facial features, reflecting specialization for speech perception. Conversely, MTG employs a feature-wide but frequency-focused coding strategy: regardless of modality, its multisensory gains converge on specific frequency windows, predominantly beta1 with modest alpha and low gamma contributions, while within these bands, it broadly integrates diverse facial action units and articulatory features. This complementary architecture of feature selectivity in STG versus frequency selectivity in MTG enables the auditory cortex to support hierarchical levels of multisensory processing, from phonetically precise speech analysis in STG to socially holistic face voice integration in MTG.

### Visual cues selectively enhance MTG-driven neural decoding and linguistic reconstruction

Finally, we asked whether these representational differences translate into functional gains for speech decoding and reconstruction. Using a dual-pathway neural speech reconstruction framework [32], we compared AV versus A decoding from STG, MTG, and combined regions (**Fig. 6, Supplementary Fig. 2**). This framework processes neural signals in parallel: an acoustic pathway reconstructs continuous acoustic features (e.g., mel-spectrograms), while a linguistic pathway decodes discrete linguistic units (e.g., phonemes or character sequences), and finally fuses the outputs to generate the speech waveform. (See Methods for details.)

**Fig. 6.**
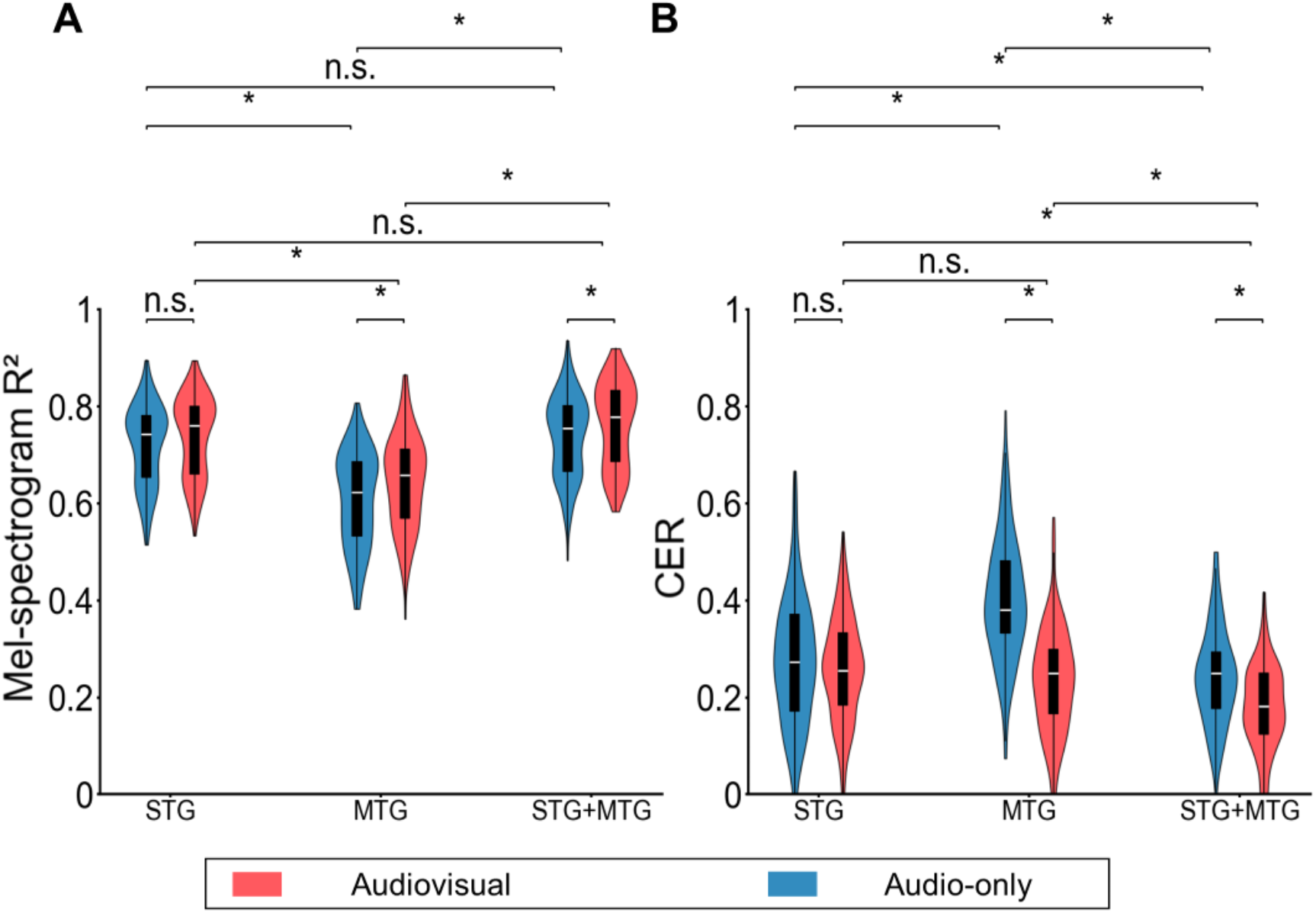
Decoding performance of audiovisual (AV) and audio-only speech from neural activity in the superior temporal gyrus (STG), middle temporal gyrus (MTG), and their combined region (STG + MTG). **A**. Mel-spectrogram correlation (R^2^) shows significant improvements for audiovisual compared to A conditions across all regions (all p < 0.01), demonstrating enhanced acoustic feature encoding with visual input. **B**. Character error rate (CER) indicates that audiovisual input leads to significantly lower error rates than audio-only input in most comparisons. Specifically, under audiovisual conditions, STG and MTG performance is comparable, while under A conditions, MTG exhibits significantly higher CER than STG. In the STG, audiovisual input significantly reduces CER compared to audio-only input, whereas in MTG, the difference between audiovisual and A is less consistently significant. Error bars represent standard deviation; * denotes statistical significance at p < 0.01.

The results revealed distinct regional dependencies on visual input. In terms of acoustic fidelity (mel-spectrogram R^2^, **Fig, 6A, Supplementary Fig. 3**), the STG exhibited remarkable stability: performance was comparable between A (R^2^ = 0.721 ± 0.082) and AV (R^2^ = 0.739 ± 0.082) conditions, and no significant difference was found (t(302) = −1.90, p = 0.0584). In contrast, the MTG showed a significant benefit from visual information, with R^2^ increasing from 0.609 ± 0.097 in A to 0.646 ± 0.095 in AV (t(302) = −3.37, p = 8.48×10^−4^). When signals from both regions were combined (STG+MTG), the AV condition also yielded a higher R^2^ (0.764 ± 0.086) compared to the A condition (0.738 ± 0.083).

For linguistic intelligibility (character error rate, CER), a similar dissociation was observed (**Fig, 6B, Supplementary Fig. 4**). The STG maintained robust decoding regardless of visual input, with CER changing only slightly from 0.277 ± 0.130 (A) to 0.260 ± 0.105 (AV) with no statistically significant difference (t(302) = 1.234, p = 0.218). The MTG, however, strongly depended on visual cues: under A alone, its CER was substantially higher (0.401 ± 0.125) than that of the STG, but with AV input, it dropped sharply to 0.234 ± 0.106 (A vs. AV: t(302) = 12.512, p = 3.19×10^−29^), even matching the performance of STG (t(302)=2.12, p = 0.070). The combined STG+MTG region achieved the lowest overall CER in the AV condition (0.182 ± 0.085), compared to 0.242 ± 0.099 in the A condition.

In summary, the STG serves as an auditory-dominant brain area for speech decoding, largely independent of visual context. The MTG plays a complementary, modality-dependent role, requiring visual input to achieve efficient linguistic decoding. Together, these regions enable robust multisensory speech reconstruction, with the combined signals offering the highest performance when both auditory and visual information are available.

## Discussion

On the basis of the previous works on speech perception, this study extends the investigation to multimodal contexts by combining naturalistic stimuli, feature-resolved encoding models, and neural decoding to provide a comprehensive, frequency-specific account of how the human temporal lobe processes auditory-visual speech. Our findings reveal that audiovisual integration in the superior temporal gyrus and middle temporal gyrus is not a monolithic process but is instead implemented through complementary, regionally specialized and frequency-dependent coding regimes that directly address several unresolved questions raised in the introduction.

A central question in the field has been whether regions traditionally associated with auditory processing (STG) and facial motion processing (MTG/STS) play distinct roles in multisensory speech[8, 11, 21, 22]. Our results provide direct evidence for a functional dissociation wherein STG employs a feature-focused integration strategy while MTG adopts a frequency-focused one. In STG, multisensory integration is characterized by selective gains for lip-reading-related facial features: the addition of visual information (AV > A) enhanced encoding of lip-related AUs in alpha and high-gamma bands but left AKT encoding largely unchanged (**Fig. 4C**; **Fig. 5C**). This suggests that STG, while primarily an auditory processor [14, 15], uses visual input to refine specific phonetic representations, aligning with recent work showing that lip-reading information penetrates auditory cortex [27, 30]. Conversely, when auditory information was added (AV > V), STG exhibited broad, multi-band enhancement of articulatory features (**Fig. 5D**), underscoring its auditory-centric nature where vision serves a modulatory rather than driving function[30]. In contrast, MTG demonstrated a frequency-focused strategy: regardless of whether the added modality was auditory or visual, multisensory gains converged predominantly on the beta1 (12–24 Hz) band, while within this band integration was feature-wide, encompassing both AUs and AKTs (**Fig. 4E-F**; **Fig. 5E-F**). This pattern supports the view of MTG as a higher-order multisensory hub [8, 9, 23], but with a critical refinement: its integrative capacity is channeled through specific spectral channels, with beta-band modulation potentially reflecting its role in maintaining sensorimotor predictions during natural communication.

Our frequency-resolved analyses also shed light on a second question raised in the introduction: how different oscillatory regimes contribute to audiovisual processing [19, 24]. The data support a division of labor across frequencies wherein low-frequency bands subserve temporal alignment while high-frequency bands support feature extraction. Low-frequency bands (Delta/Theta) showed enhanced encoding of temporal dynamics, particularly for articulatory kinematics in MTG under AV conditions (**Fig. 5B**), likely reflecting a mechanism for aligning the temporal structure of auditory and visual streams to enable multisensory coherence [19]. High-gamma activity, by contrast, was strongly engaged in STG during encoding of fine-grained articulatory detail and lip movements (**Fig. 4A, 5A**), consistent with its established role in local cortical computation and feature extraction [14, 16, 26]. Notably, the beta band (12–24 Hz) emerged as a key nexus for integration in MTG, bridging the gap between low-frequency tracking and high-frequency feature encoding—a finding that resonates with work implicating beta oscillations in maintaining sensory predictions and coordinating top-down and bottom-up signals during speech perception[31].

Finally, our acoustic and lexical dual-pathway decoding results provide functional validation of these encoding principles by demonstrating that visual information confers significant benefits for speech reconstruction, particularly for MTG and for linguistic intelligibility (**Fig. 6**). The STG maintained robust decoding performance even without visual input, reinforcing its role as an auditory-anchored processor consistent with its established role in representing spectrotemporal acoustics and phonetic features[12, 13]. However, the MTG’s decoding accuracy was heavily dependent on visual cues; without them its character error rate (CER) was substantially higher, but with them it matched or even exceeded STG performance. This suggests a functional hierarchy wherein STG provides a stable, modality-general representation of speech acoustics while MTG leverages visual input to resolve linguistic ambiguity and enhance semantic clarity, likely by accessing a richer representation of the speaker’s communicative actions [5, 6, 24]. The combined STG+MTG decoder achieving the highest performance indicates that these regions contribute non-redundant information for robust speech perception.

While our study fundamentally addresses the perceptual integration of speech, the frequency-specific and cross-modal architectures identified here offer crucial computational insights for the development of speech brain-computer interfaces (BCIs). Current speech neuroprostheses primarily target motor and premotor cortices to decode intended articulation. However, our findings demonstrate that robust decoding of naturalistic speech relies heavily on multiplexing low-frequency kinematic tracking with high-frequency feature extraction. This frequency-resolved computational strategy provides a highly transferable blueprint for BCI decoding algorithms[33]. By successfully applying a dual-pathway acoustic and linguistic framework to continuous Mandarin, we demonstrate how integrating distinct sensory-kinematic priors (such as AKTs and AUs) can drastically reduce decoding errors[34–36]. Moving forward, adopting these multimodal, cross-frequency decoding strategies will be instrumental in developing next-generation neuroprostheses capable of highly accurate, context-aware communication, particularly for the complex neural dynamics inherent to tonal languages[37, 38].

While our work provides a detailed account of cortical surface processing, it is limited by the inability to directly record from the superior temporal sulcus (STS), a deep structure critical for multisensory binding [10, 11, 28], future studies using stereo-EEG will be essential to probe the role of the STS in coordinating the cross-frequency dynamics observed here. Furthermore, our passive viewing paradigm could be extended to include active tasks or audiovisual conflict (e.g., McGurk stimuli) to dissociate automatic integration from perceptual decision-making[1, 7].

In summary, this study reveals that the human temporal lobe employs complementary coding strategies for audiovisual speech: STG implements a feature-selective, auditory-dominant regime that uses visual lip cues to refine phonetic representations, while MTG implements a frequency-selective regime that uses the beta band as a hub to broadly integrate facial and articulatory features, supporting context-sensitive perceptual synthesis. Together, these mechanisms enable robust speech understanding in complex, naturalistic settings. Ultimately, the frequency- and region-specific principles we have identified could inform next-generation brain-computer interfaces that leverage low-frequency signals for rhythmic alignment and high-frequency signals for articulatory detail to achieve robust, multimodal speech decoding in real-world environments.

## Methods

### Participants

A total of 8 participants (4 males, 4 females) were recruited for this study. The cohort included 6 individuals with eloquent brain tumors, who underwent intraoperative awake surgery following routine clinical protocols, and 2 patients with epilepsy who were undergoing comprehensive presurgical assessment to determine optimal surgical strategies. Each participant provided informed consent prior to surgery. At that time, it was clearly explained, as outlined in the IRB-approved written protocol and consent document, that the research task was for scientific purposes only and would not directly affect their clinical care. It was also emphasized to each participant that their involvement in the research task was entirely voluntary. All procedures were approved by the Institutional Review Board of Huashan Hospital, Fudan University (brain tumor partcipants: approval no. KY2024-842; epilepsy patients: approval no. KY2023-883).

### Data acquisition and neural signal processing

For intraoperative paticipants, one or two 128-channel ECoG grids with 4 mm center-to-center spacing and 1.17 mm contact diameter were temporally placed on the cortical surface. Signals were recorded at 3052 Hz using a Tucker-Davis Technologies system. For epilepsy patients, synchronized ECoG and audio signals were acquired at 15 kHz using an implanted neural interface (Neuroxess Co. Ltd). The integrated implant consisted of a 256-channel flexible high-density ECoG array (1.3 mm contact diameter, 3 mm interelectrode spacing), a flexible printed circuit for signal transmission, a signal processing unit with four Intan RHD2164 chips for noise suppression and amplification, and a titanium enclosure ensuring waterproofing and biocompatibility for stable long-term operation. All neural signals underwent consistent processing across seven frequency bands: Delta (0.5-4 Hz), Theta (4-8 Hz), Alpha (8-12 Hz), Beta1 (12-24 Hz), Beta2 (24-40 Hz), Low Gamma (40-70 Hz), and High Gamma (70-150 Hz). Each band was processed independently through the following steps: bad channel rejection, common average referencing, band-specific filtering, analytic amplitude extraction via Hilbert transform, and z-score normalization per channel across time. For offline analysis, signals were downsampled to 400 Hz. Line noise and harmonics were removed using notch filters adapted for the recording environment.

### Experimental Stimuli

The experimental stimuli were constructed using video materials from the Chinese Mandarin Lip Reading (CMLR) dataset[39] (available at https://vipazoo.cn/CMLR.html). The stimulus set consisted of 40 complete paragraphs, comprising 281 sentences spoken by 11 professional news anchors (7 male, 4 female). Each paragraph contained 4-12 sentences forming a coherent discourse unit, with paragraphs separated by 0.5-second intervals while sentences within paragraphs were presented continuously.

The experiment comprised nine blocks with the following sequence: audiovisual block 1 (AV1), audio-only block 3 (A3), audiovisual block 2 (AV2), audio-only block 4 (A4), video-only block (V), audio-only block 1 (A1), audiovisual block 3 (AV3), audio-only block 2 (A2), and audiovisual block 4 (AV4). This design included four audiovisual blocks, four audio-only blocks (using the same auditory materials as corresponding AV blocks but without visual components), and one video-only block featuring the initial sentences from all 40 paragraphs. Each block contained 10 paragraphs and had a maximum duration of 5 minutes.

All stimuli featured professional news anchors maintaining stable head position in the studio setting, with synchronized articulatory movements and natural speech delivery. The actual number of blocks completed by each patient is detailed in Supplementary Table 1.

### Electrode localization

In cases involving chronic monitoring, the electrode placement procedure involved pre-implantation MRI and post-implantation CT scans. In awake cases, temporary high-density electrode grids were utilized to capture cortical local potentials, with their positions recorded using the Medtronic neuronavigation system. The recorded positions were then aligned to pre-surgery MRI data, with intraoperative photographs serving as additional references. Localization of the remaining electrodes was achieved through interpolation and extrapolation techniques.

### Facial Action Units (AUs), Articulatory Kinematic Trajectories (AKT) and their feature extractions

Facial Action Units (AUs) are the fundamental components of the Facial Action Coding System (FACS), which describes facial expressions based on the contraction of specific facial muscles or muscle groups. Each AU corresponds to a minimal, anatomically-based action, such as inner brow raiser (AU1) or lip corner puller (AU12). To quantify facial behavior in naturalistic settings, we employed automated facial behavior analysis software, OpenFace (v2.0), a state-of-the-art computer vision tool capable of detecting facial landmarks, estimating head pose, and recognizing the presence and intensity of AUs from standard 2D video recordings. For each video frame, OpenFace outputs a vector of continuous intensity scores and detection probabilities (0/1) for a predefined set of AUs. This approach provides a non-invasive, model-free description of facial kinematics, analogous to the use of acoustic-to-articulatory inversion for inferring vocal tract movements, as it translates readily observable visual signals (video pixels) into a standardized set of physiologically meaningful movement descriptors without requiring direct muscle or motion capture.

Articulatory Kinematic Trajectories (AKTs) represent the coordinated, time-varying movements of vocal tract articulators (e.g., tongue, lips, jaw) that underlie continuous speech production, as defined in the seminal work by Chartier et al.. The core methodology for obtaining AKTs involves a speaker-independent Acoustic-to-Articulatory Inversion (AAI) model. This deep learning-based approach infers the kinematic trajectories of articulators from speech acoustics alone, without requiring direct articulatory measurements from the target speaker. The model was trained on a multi-speaker database containing simultaneous recordings of speech audio and Electromagnetic Midsagittal Articulography (EMA) data, which provides ground-truth Cartesian coordinates of articulator movements. The AAI model maps acoustic features (e.g., mel-cepstral coefficients) combined with phonological context to the kinematic space. The resulting inferred trajectories, capture the coordinated, often “out-and-back,” dynamic patterns of multiple articulators working in concert to achieve specific vocal tract configurations (e.g., coronal or labial constrictions).

The extraction of AUs and AKTs follows a conceptually parallel pipeline, transforming raw, naturalistic behavioral signals into interpretable kinematic features. For AUs, the input is a 2D video stream. The process involves face detection, landmark localization, and regression/classification to map facial appearance features onto AU intensities. For AKTs, the input is an acoustic speech signal. The process involves acoustic feature extraction, followed by a deep neural network mapping to an articulatory kinematic space. Both methods are inversion problems: they infer underlying, unobserved physiological kinematics (facial muscle movements or vocal tract articulator movements) from their readily recorded external manifestations (video or audio). This allows for the study of complex, continuous natural behavior without intrusive instrumentation. A key parallel finding is that both AUs and AKTs often represent coordinated movement patterns rather than isolated actions, evidenced by the multi-articulator nature of AKTs and the co-occurrence patterns of AUs in facial expressions.

### Responsive electrode selection undergoing audiovisual, audio-only and video-only stimuli

To identify electrodes responsive to audiovisual (AV), audio-only (A), and video-only (V) stimuli, we analyzed recorded neural activity aligned to stimulus onset. For each electrode and each stimulus condition, we compared the post-stimulus time course (from 0 ms onward) to a resting baseline defined as the interval from −400 to −200 ms relative to stimulus onset. An electrode was considered responsive in a given modality if there existed at least one continuous 200 ms time window during which every individual time point showed an ERP response significantly different from the rest-state baseline (p < 0.01, two-sided t-test, Bonferroni-corrected across all tested electrodes). This criterion ensured that only electrodes exhibiting sustained, statistically reliable modulation by the stimulus, rather than transient fluctuations, were included in subsequent analyses.

### Temporal receptive field (TRF) analysis and Unique R^2^ analysis

To quantify how auditory and visual stimuli dynamically modulate neural activity across time, we employed temporal receptive field (TRF) analysis, a model-based approach that estimates the linear relationship between time-varying input features and neural responses. For each participant, TRF models were trained using ridge regression with a fixed temporal window of −400 to 0 ms relative to stimulus onset, divided into 20 ms bins. The TRF weights, representing the contribution of each time bin to the neural response, were normalized by the maximum absolute value within the window to ensure comparability across conditions. Cross-validation (leave-one-trial-out) was used to optimize hyperparameters (regularization strength λ) and assess model generalization. TRF results were then projected onto cortical surfaces for visualization, with statistical significance determined via cluster-based permutation tests (p < 0.05, FDR-corrected).

To disentangle the independent contributions of articulatory kinematic trajectories (AKTs) and facial action units (AUs) to neural encoding, we applied unique R^2^ analysis, a multivariate method that isolates the variance uniquely explained by each feature group while controlling for overlapping contributions from other inputs. Specifically, we constructed full models including all feature groups (e.g., visible articulators, larynx, AUs) and reduced models excluding the target feature group. For each condition (audio-only [A], video-only [V], audiovisual [AV]), the unique R^2^ of a specific feature group was calculated as the difference between the total R^2^ of the full model and the R^2^ of the reduced model (i.e., 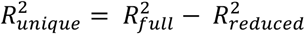). This process was repeated across frequency bands (delta: 0.5–4 Hz; theta: 4–8 Hz; alpha: 8–12 Hz; beta1: 12–24 Hz; low gamma: 40–70 Hz) using bandpass-filtered data. Statistical comparisons between conditions (e.g., AV > A, AV > V) were performed with non-parametric permutation tests (1,000 iterations), with multiple comparisons corrected using false discovery rate (FDR) control.

These complementary methods allowed us to systematically evaluate how temporal dynamics (via TRF) and modality-specific contributions (via unique R^2^) shape neural representations of speech-related and expressive facial features, as illustrated in Fig. 4 and 5.

### The dual-pathway neural speech reconstruction framework employed for decoding speech from neural activity

To investigate the encoding of speech by neural populations recorded from the STG and MTG from a decoding perspective, we utilized the following dual-pathway neural speech reconstruction framework. This framework enables the analysis of how information is represented within these regions by attempting to reconstruct the original speech signal from the neural activity patterns.

The framework consists of two main pathways operating in parallel. The acoustic pathway takes neural signals as input and aims to reconstruct continuous acoustic features, specifically mel-spectrograms. It utilizes an acoustic feature generator, implemented using an RVQGAN-based decoder architecture[40]. RVQGAN (Vector Quantized Generative Adversarial Network) is effective in generating high-fidelity acoustic representations from latent neural codes. The linguistic pathway processes the same neural signals to decode the underlying linguistic content, such as phoneme sequences or character transcripts. The output from this pathway serves as conditioning information for the subsequent synthesis step. The linguistic pathway employs a Text-to-Speech (TTS) generator, specifically adapted from the CosyVoice 2.0 model[41]. CosyVoice 2.0, known for its capabilities in voice cloning and text-to-speech synthesis, provides robust linguistic feature extraction and can synthesize speech based on the decoded linguistic units and acoustic priors.

Finally, the outputs from both pathways (the reconstructed mel-spectrogram from the acoustic pathway and the linguistic representation/synthesis guidance from the linguistic pathway) are integrated to produce the final reconstructed speech waveform using a CosyVoice 2.0 voice cloner. This dual-pathway design allows for the independent modeling of acoustic fidelity and linguistic content, integrating the strengths of specialized models for each aspect before combining them for the ultimate goal of speech synthesis.

### The training detail of the employed dual-pathway framework

To achieve high-fidelity speech reconstruction from the limited clinical neural data, both pathways in the framework were trained under a unified strategy of alignment to pre-trained deep speech generators with lightweight adaptation. The core principle involved keeping the parameters of the powerful, pre-trained backbone models (the RVQGAN-based acoustic decoder and the CosyVoice 2.0 TTS/cloner) completely frozen. Only the newly introduced, light-weight neural adapters were trained to learn the mapping from the neural signals to the respective pre-defined latent spaces. This approach maximizes data efficiency and prevents overfitting.

For each participant, approximately 15 minutes of temporally aligned neural recordings (ECoG signals) and their corresponding speech stimuli (audio waveforms) were used. This paired dataset was strictly partitioned into training, validation, and test sets with a ratio of 34:3:3. This ensured sufficient data for model learning while reserving independent sets for hyperparameter tuning and final unbiased evaluation.

The acoustic adapter was tasked with mapping the multi-band neural features to a 1024-dimensional acoustic latent space, which serves as the input to the frozen RVQGAN-based acoustic speech decoder. The training objective was to minimize the difference between the predicted and the target acoustic features (extracted from the ground-truth speech by the frozen feature extractor). This was achieved using a Sigmoid-scaled Mean Squared Error (MSE) loss applied to the feature vectors.

The AdamW optimizer with its default parameters (learning rate lr=10^−3^, *β*_1_=0.9, *β*_2_=0.999) was employed. The model was trained for 200 epochs, a process taking approximately 4 hours on a single NVIDIA RTX 3090 GPU. Training was monitored using the validation loss, with early stopping implemented to prevent overfitting.

The linguistic adapter, based on a Transformer architecture, was trained to decode neural activity into a sequence of 512-dimensional semantic feature vectors. Its composite loss function combined two terms: A Sigmoid-scaled MSE feature loss to ensure semantic similarity, and a 0.01-weighted length prediction loss to encourage accurate sequence length estimation, aiding in structured generation. Optimization was performed using the default AdamW optimizer. Training also proceeded for 200 epochs (~4 hours), with the character error rate (CER) on the validation set guiding model selection.

The two adapters were trained independently. After training, the complete decoding pipeline operates as follows: neural signals are fed concurrently into both trained adapters. The acoustic adapter’s output drives the RVQGAN decoder to produce a mel-spectrogram rich in speaker-dependent acoustic details. Simultaneously, the linguistic adapter’s output provides the semantic and textual guidance to the CosyVoice 2.0 voice cloner. Finally, the cloner integrates the generated mel-spectrogram (acoustic blueprint) and the linguistic instructions to synthesize the final, high-quality speech waveform that balances naturalness and intelligibility.

## Data availability

The data that support the findings of this study are available on request from the lead contact. The data are not publicly available because they could compromise research participant privacy and consent.

## Code availability

All original code and preprocessed anonymized data to replicate the main findings of this study can be found at https://github.com/CCTN-BCI/Audiovisual_integration. Any additional information required to reanalyze the data reported in this paper is available from the lead contact upon request.

## Acknowledgments

This work is supported by the Brain Science and Brain-like Intelligence Techonology - National Science and Technology Major Project of China (2025ZD0217000 to Y.L. and 2022ZD0212300 to J.Lu), the National Natural Science Foundation of China (32371154 to Y.L. and 32371146 to J.Lu), Shanghai Rising-Star Program (24QA2705500 to Y.L.), the Lin Gang Laboratory (LG-GG-202402-06 and LGL-1987-18 to Y.L.), the Innovation Program of Shanghai Municipal Education Commission (2023ZKZD13 to J.W.), and the Shanghai Oriental Talent Program Leading Project (DFYCLJ04 to J.W.). The computations in this research are supported by the HPC Platform of ShanghaiTech University.

## Author contributions

Conceptualization: Y.L. and J. Lu; Methodology: J. Li, K.B. and Y.L.; Software: J.Li and Y.L.; Formal analysis: J. Li, K.B., X.H., J. Lu and Y.L.; Resources: J.W., J. Lu, and Y.L.; Writing— original draft: J. Li; Writing—review and editing: J.Li, J. Lu and Y.L.; Supervision: J. Lu and Y.L.; Project administration: Y.L.

