## Supplementary Information for "Dissociable frequency regimes in human temporal cortex integrate facial and acoustic cues during natural speech"

Li *et al*

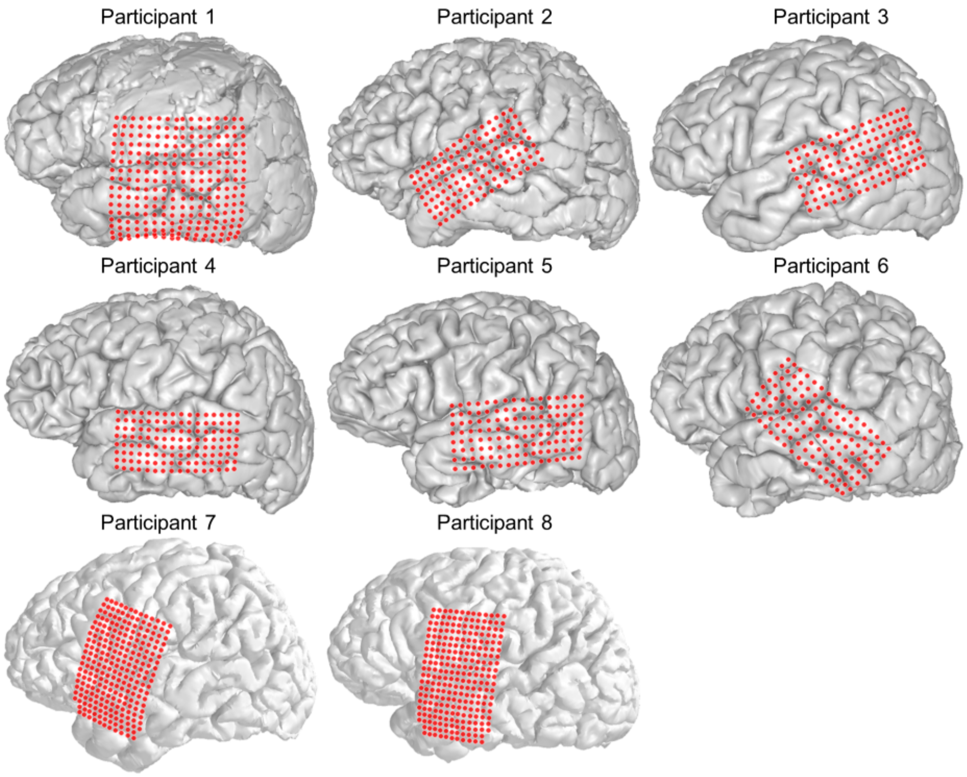


**Supplementary Figure 1. Electrodes for all participants.** The locations of the ECoG electrodes were plotted with red scatters.

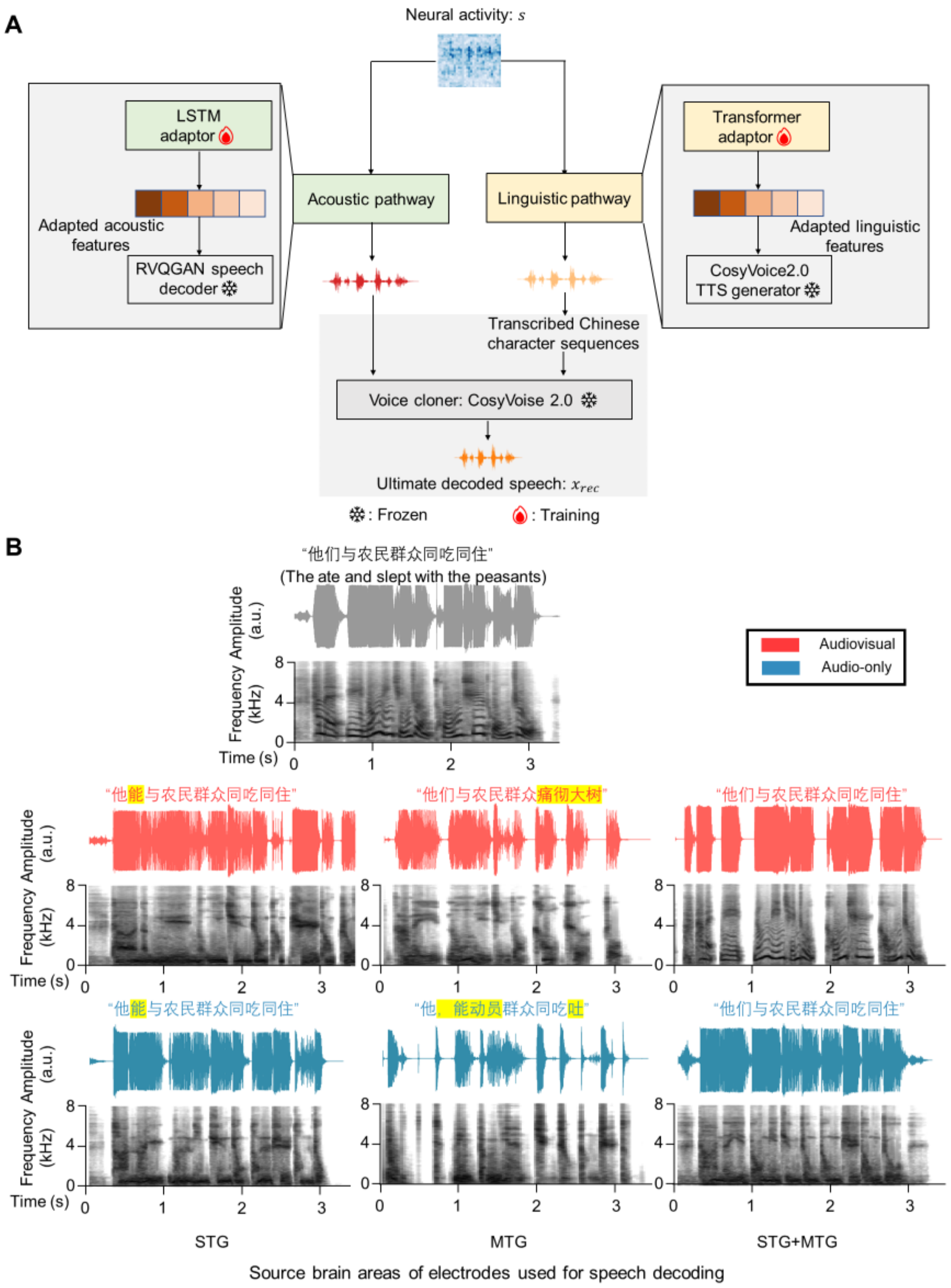


**Supplementary Figure 2. The architecture of the acoustic-linguistic dual-pathway framework for neural-driven Mandarin speech reconstruction and demonstrative speech waveforms.** (**A**) The framework architecture and training protocol. The system consists of an acoustic pathway and a linguistic pathway, both driven by neural activity. In the acoustic pathway, a lightweight LSTM adaptor (marked with a red dot for training) maps neural features to a 1024-dimensional acoustic latent space, which is fed into a frozen RVQGAN speech decoder to generate high acoustic fidelity speech. In the linguistic pathway, a Transformer adaptor (marked with a red dot for training) maps neural features to 512-dimensional semantic vectors, which are processed by a frozen

CosyVoice 2.0 TTS generator to produce transcribed Chinese character sequences. Both pathways converge at the final voice cloning stage, where a frozen CosyVoice 2.0 model integrates the acoustic blueprint and linguistic content to synthesize the ultimate decoded speech.

(**B**) Decoded speech examples from different brain areas. The top panel shows the ground truth audiovisual speech waveform and its corresponding mel-spectrogram for the sentence "世界多极化经济全球化进一步发展" (The trend towards a multipolar world and economic globalization is further developing). The middle and bottom panels display the reconstructed speech waveforms and mel-spectrograms for audiovisual (red) and audio-only (blue) conditions, respectively, decoded from neural signals recorded in the superior temporal gyrus (STG), middle temporal gyrus (MTG), and the combined STG+MTG regions. The decoded text snippets below each reconstruction highlight variations in linguistic accuracy across conditions and brain areas.

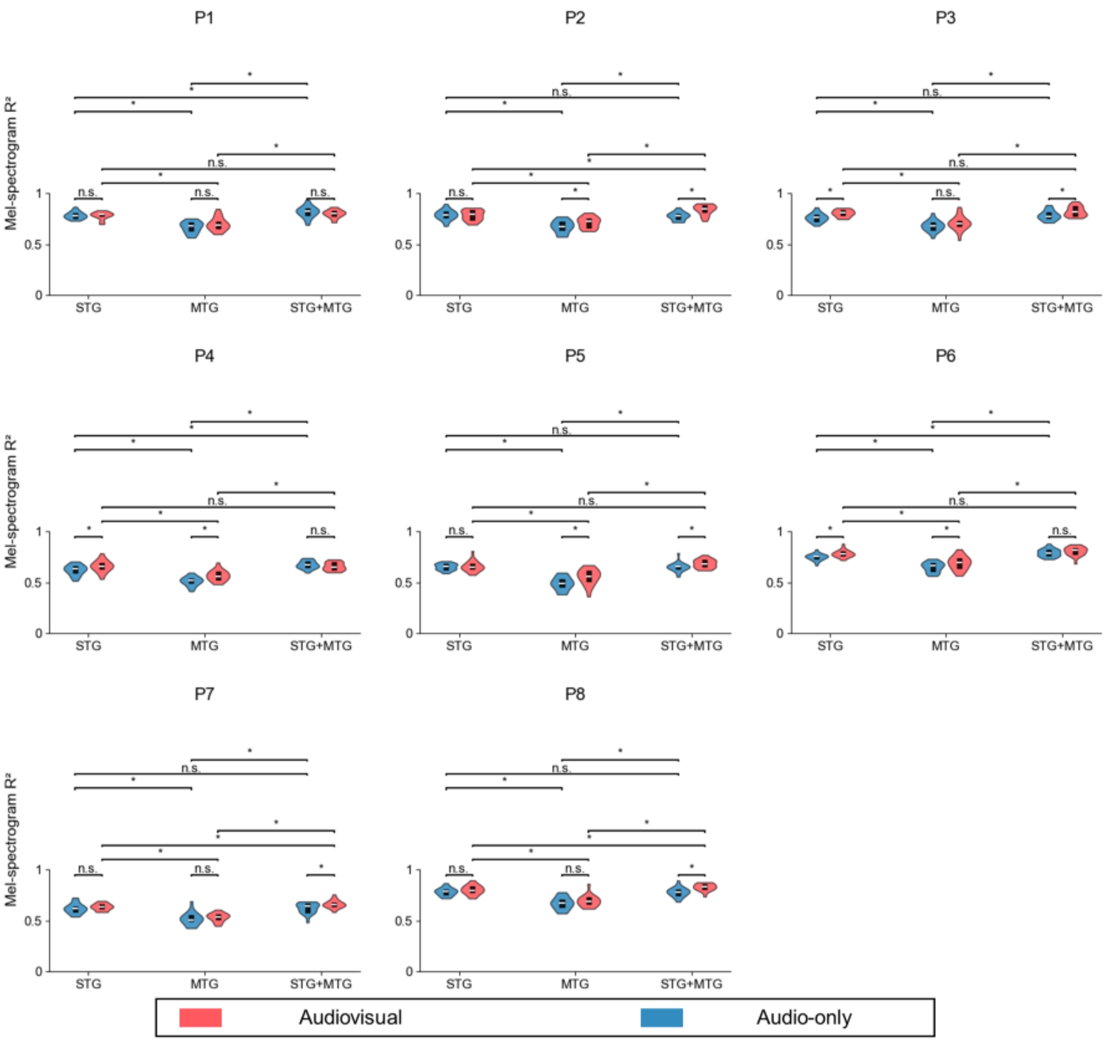


**Supplementary Figure 3. Individual participant-level analysis of mel-spectrogram reconstruction $R^2$ for audiovisual and audio-only conditions across the STG, MTG, and STG+MTG**. Each panel shows violin plots representing the distribution of $R^2$ for one participant. Red indicates audiovisual condition; blue indicates audio-only condition. Statistical significance is denoted by * ($p < 0.05$) or n.s. (not significant) based on paired comparisons within each participant and region.

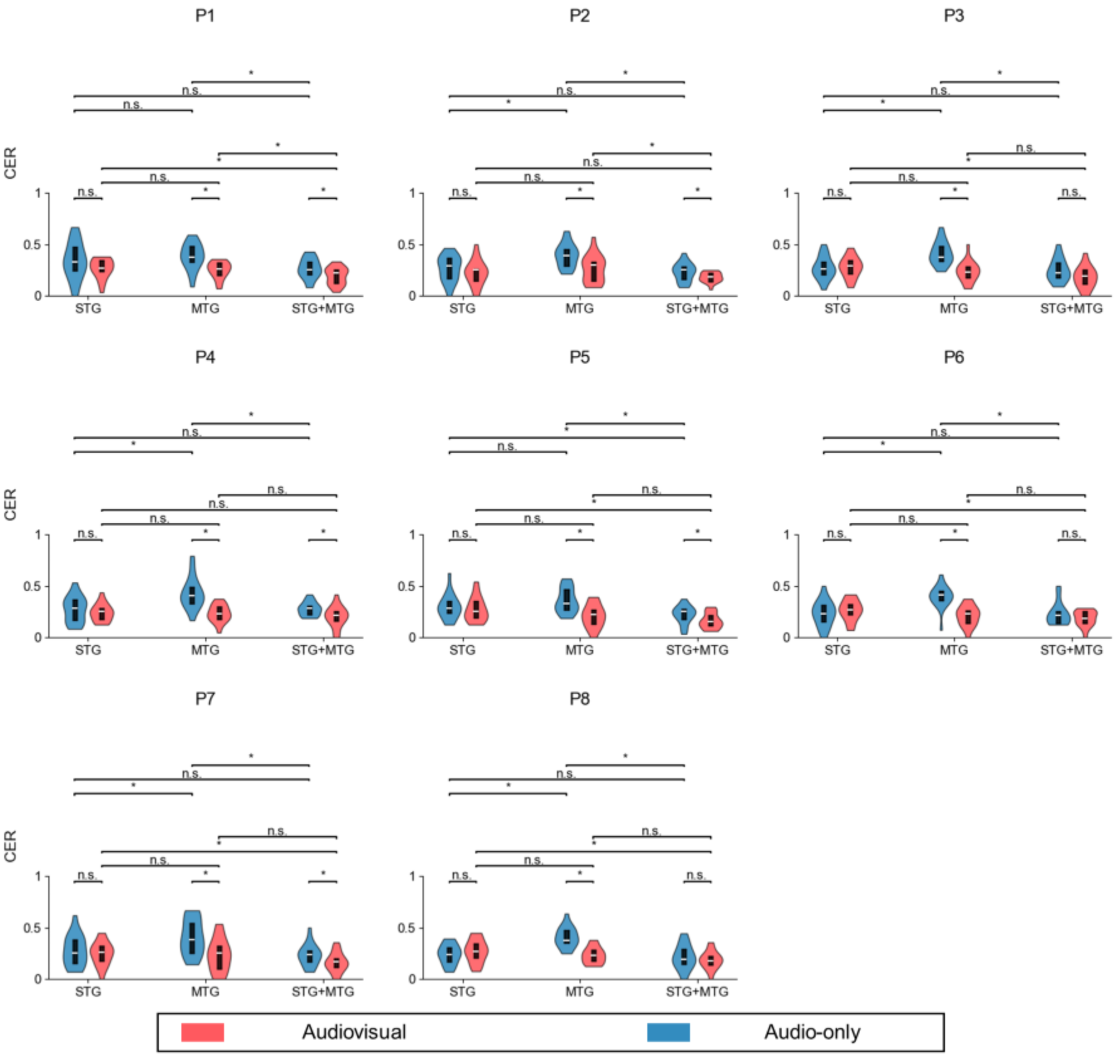


**Supplementary Figure 4. Individual participant-level analysis character error rate (CER) for audiovisual and audio-only conditions across the STG, MTG, and STG+MTG**. Each panel shows violin plots representing the CER for one participant. Red indicates audiovisual condition; blue indicates audio-only condition. Statistical significance is denoted by * ($p < 0.05$) or n.s. (not significant) based on paired comparisons within each participant and region.
